# Comparison of RNA-seq and Microarray Platforms for Splice Event Detection using a Cross-Platform Algorithm

**DOI:** 10.1101/197798

**Authors:** Juan P. Romero, María Ortiz-Estévez, Ander Muniategui, Soraya Carrancio, Fernando J. de Miguel, Fernando Carazo, Luis M Montuenga, Remco Loos, Rubén Pío, Matthew W. B. Trotter, Angel Rubio

**Affiliations:** CEIT and Tecnun, University of Navarra, San Sebastián, Spain; Celgene Institute for Translational Research Europe, Celgene Corporation, Seville, Spain; Program in Solid Tumors and Biomarkers, CIMA, University of Navarra, Pamplona, Spain; Department of Histology and Pathology, University of Navarra, Pamplona, Spain; IdiSNA, Navarra Institute for Health Research, Pamplona, Spain; Department of Biochemistry and Genetics, University of Navarra, Pamplona, Spain; CIBERONC, Centro de Investigación Biomédica en Red, Madrid, Spain

**Author notes:** Corresponding author: Angel Rubio, CEIT and TECNUN, University of Navarra, Manuel Lardizabal 13-15, San Sebastian, Spain. Join first authors (Juan P. Romero, María Ortiz-Estévez).

**Keywords:** Alternative splicing, RNA-seq, Microarrays

## Abstract

RNA-seq is a reference technology for determining alternative splicing at genome-wide level. Exon arrays remain widely used for the analysis of gene expression, but show poor validation rate with regard to splicing events. Commercial arrays that include probes within exon junctions have been developed in order to overcome this problem.

We compare the performance of RNA-seq (Illumina HiSeq) and junction arrays (Affymetrix Human Transcriptome array) for the analysis of transcript splicing events. Three different breast cancer cell lines were treated with CX-4945, a drug that severely affects splicing. To enable a direct comparison of the two platforms, we adapted EventPointer, an algorithm that detects and labels alternative splicing events using junction arrays, to work also on RNA-seq data. Common results and discrepancies between the technologies were validated and/or resolved by over 200 PCR experiments.

As might be expected, RNA-seq appears superior in cases where the technologies disagree, and is able to discover novel splicing events beyond the limitations of physical probe-sets. We observe a high degree of coherence between the two technologies, however, with correlation of EventPointer results over 0.90. Through decimation, the detection power of the junction arrays is equivalent to RNA-seq with up to 60 million reads. Our results suggest, therefore, that exon-junction arrays are a viable alternative to RNA-seq for detection of alternative splicing events when focusing on well-described transcriptional regions.

## Introduction

Alternative Splicing (AS) is known to play a major role in human biology, and the identification of transcriptional splicing patterns has potential uses for diagnosis, prognosis, and therapeutic target evaluation in the disease context [1,2]. The development of exon microarrays enabled the transcriptomic study of differential splicing events, but PCR validation rates for identification of splice differences via microarray analysis tend to be lower than those observed for identification of differential gene expression using similar technologies [3–5]. Junction arrays [6–10] have been proposed to overcome this problem by using oligonucleotide probe-sets that interrogate junctions between exons in the transcriptome, as well as the exons themselves.

Since the advent of next-generation sequencing (NGS), RNA-seq has become the technology of choice via which to detect and quantify alternative splicing (for a review see [11]). Various published works compare the performance of RNA-seq and expression microarrays for the analysis of gene expression [12,13], but a thorough evaluation of both technologies in terms of their ability to detect differential AS events has yet to be presented. In the present study, we perform a comparison of RNA-seq technology (using the Illumina HiSeq platform) and junction arrays commercialized by Affymetrix (Human Transcriptome array, or HTA).

AS can be studied from two complementary points of view: with focus on transcripts or splicing events respectively. In the former, the subject of analysis is the transcript (or isoform), whereas in the latter, the subject(s) are the splicing events themselves.

The pipeline of the transcript-focused approach uses RNA-seq data with and without known annotations in order to reconstruct the transcriptome and estimate the concentration values of the transcripts. Finally, the significance of change in absolute or relative concentrations is assessed using suitable statistical methods [14–16]. Transcript reconstruction is challenging [17] and any error in reconstruction of transcript structure may be propagated to the estimation of corresponding concentrations and thereby to the output of statistical analysis. Moreover, the challenge of estimating isoform concentrations for genes with many transcripts yields wide confidence intervals[18].

In order to circumvent the problem of transcript reconstruction, the transcriptome may be taken as algorithmic input in RNA-seq when performing a direct estimation of isoform concentrations [19]. This approach, however, only measures annotated transcripts and is hence unable to detect and quantify novel isoforms. Despite this obstacle, methods to estimate isoform concentrations using microarrays have been proposed [20–23]. A recent comparison of the isoform deconvolution using both RNA-seq and microarrays has been published using non-junction Affymetrix HuEx arrays [24].

One potential criticism of transcript-focused methods is that they may miss local AS variability because of the inherent challenge of isoform deconvolution [25]. Even the better methods display transcriptome reconstruction levels below 50% when using simulated reads, i.e. less than 50% of transcripts are recovered and less than 50% of predictions are correct [17]. On this basis, therefore, an event-based method appears a more suitable approach via which to compare AS detection technologies, with the additional benefit of straightforward validation using PCR.

Event-based methods focus directly on the analysis of differential splicing events, rather than first attempting to estimate transcript concentration levels. These events can be classified into five canonical categories[26]: cassette exon, alternative 3′, alternative 5′, mutually exclusive exons and intron retention. In some cases, alternative start and termination sites are included also when defining splicing events. This approach has gained traction and several algorithms have been developed recently for detection of splicing events using RNA-seq data, including rMats, SplAdder, spliceGrapher or SGSeq [27–30]. SpliceGrapher and SGSeq detect events prior to application of separate software in order to state corresponding statistical significance, whereas rMats and SplAdder perform both detection and statistical analysis. Alongside NGS-based approaches, AS event detection methods are available for exon arrays [31], and exon-junction arrays [6,8,9,32]. The latter methods display validation rates well above 50%.

The principal aim of this work is to compare RNA-seq and exon-junction microarray technologies in their ability to detect differential AS events. To do so comprehensively, and to allow as close to a direct comparison as possible, we have adapted the EventPointer [8] algorithm for application to data from both platforms, generated from the same control experiment. The control experiment comprises three distinct triple-negative breast cancer (TNBC) cell-lines, exposed in culture to a drug known to affect the transcriptional machinery and, thereby, to induce AS events.

Further to comparative analysis of the resulting data, we conclude that both technologies show considerable concordance with high PCR validation rates, and that exon-junction microarrays have potential as an alternative to RNA-seq profiling for detection of AS events in annotated transcripts.

## Results

CX-4945 is a potent and selective orally bioavailable small molecule inhibitor of casein kinase CK2 [33], which has been proposed previously as a cancer therapy [34], and which has been shown to regulate splicing in mammalian cells [35]. RNA samples taken from three distinct triple-negative breast cancer (TNBC) cell-lines, exposed to CX-4945 and also to a DMSO control, were profiled using both RNA sequencing ^1^ and hybridization to exon-junction microarrays (see Methods for details). We extended the EventPointer algorithm (available via Bioconductor, see Methods and Supp. File 2) for application to data from both platforms and applied it to the corresponding datasets in order to identify AS events.

Prior to the comparison of platforms for splice event detection, the data was assessed at the gene level in order to ensure signal quality and coherence. Gene expression was computed from RNA-seq data using Kallisto [19] to quantify expression as the sum of isoform concentrations for each gene. RMA [36] was used to quantify gene expression from microarray data, using annotation files from Brainarray [37]. The same version of the Ensembl Transcriptome (Ensembl v.74, GRCh 37) was used in both cases.

Considering each technology independently, correlation between sample replicate profiles in each cell-line and experimental condition is high for both platforms (correlation coefficient ranging from 0.988 to 0.996 in arrays and 0.996 to 0.997 in RNA-seq). When comparing profiles from the same samples between technologies, strong coherence is observed for well-expressed genes. Median correlation of gene expression between technologies on the same samples is 0.510, and gene expression patterns across all samples display correlation of 0.680 between technologies. The first one is smaller owing to the different probe affinities of the set of probes that interrogates each gene. When only the 50% most highly expressed genes are considered, the median correlation of gene expression patterns is 0.750 (Supplementary file 1). The gene expression correlations observed are similar to previously reported comparisons between RNA-seq and exon arrays[38].

### Events detected by RNA-seq and junction arrays show strong qualitative and quantitative concordance, with a subset detected exclusively by one of the technologies

Figure 1 depicts the EventPointer pipeline for both profiling technologies (see original publication [8] for further detail), with CEL files (microarray) or BAM files (RNA-seq) as starting input. When building the splicing graph, each exon is split into two nodes that correspond to its start and end genomic positions respectively (Figure 1.b). Each event is described by two alternative paths (Paths 1 and 2) and a shared reference path (Path Ref) within the splicing graph. These paths are sets of edges in the splicing graph. Paths 1 and 2 are mutually exclusive in terms of isoforms (i.e. if an isoform includes Path 1 it does not include Path 2 and vice versa) and all isoforms interrogated by the event share the reference path. Therefore, events are contained in several isoforms (at least two). A simple example would be the cassette exon shown in Figure 1.b: the reference path is composed by the edge that links nodes 6a and 6b (i.e. the coverage of exon 6 or the signal in the probe-set of the array that interrogates this exon) and the edge that links nodes 8a and 8b (coverage of exon 8). All these measurements are summarized into one average value. Path 1 includes the edges in the path 6b-7a-7b-8a (coverage of exon 7 and its flanking junctions) and Path 2 is the edge that links 6b and 8a (coverage of the skipping junction). EventPointer distinguishes between events as corresponding to: cassettes; alternative 5′; alternative 3′; mutually exclusive exons; alternative first exons; and alternative end exons. Complex events that do not match any of these categories are denoted as such.

**Figure 1.**
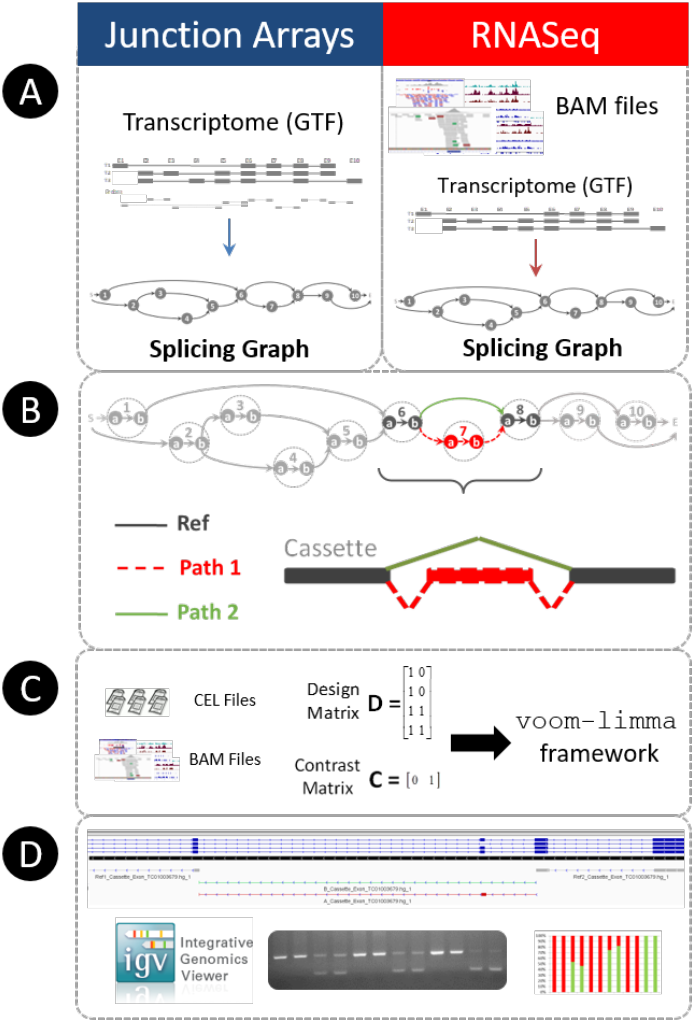
EventPointer overview for junction arrays and RNA-Seq data. A: The CEL or BAM files are the input data for each technology. The splicing graph for each gene is built using the array annotation files or directly using the sequenced reads. B: Each node in the splicing graph is splitted into two nodes that correspond to the start and end positions in the genome respectively. EventPointer identifies events within each gene and annotates the type of event. In the figure, among the events in the gene, an exon cassette is highlighted. C: Statistical significance of the events is computed. D: Finally, the top-ranked events are validated using PCR and the results visualized in IGV.

Three different cell-lines were profiled, each exposed to CX-4945 and DMSO respectively across five replicates. AS differences were tested using a linear model which controlled for cell-line differences. Using the read coverage (or probe-set signal) for each path, a statistical analysis based on voom-limma [16,39] is applied to determine the significance of each event via comparison between alternative path signals (see Material and Methods for details). In addition to the statistical analysis, we compute the Percent Splice Index (PSI or Ψ) [40], an estimate of relative isoform concentrations that map to paths 1 and 2 for each event.

In order to identify well-expressed events (more likely to be biologically significant and less prone to validation error), the comparison of AS detection was performed on a subset of the data with expression above a set threshold (see Methods for details). In brief, a junction coverage threshold was applied to the RNA-seq data (default 2 FPKM) and a threshold on expression percentile applied to the microarray data (default probe-set expression greater than 25% of probes in any sample profile).

Table 1 displays the number of significantly differentially expressed events detected via application of EventPointer to RNA-seq and microarray data respectively as these thresholds are varied. The test compares differential AS in profiles from cell-lines treated with CX-4945 and with DMSO control. As may be expected, setting more stringent expression thresholds yields fewer events detected with better False Discovery Rate (FDR) on both platforms.

**Table 1.** Number and statistical significance of detected AS events using both RNA-seq and array technologies. for different expression thresholds, default filters are junction coverage greater than 2 FPKM for RNA-seq and probe-set signal greater than top 25% quantile for microarray.

| RNA-seq |  | Exon-junction arrays |  |
| --- | --- | --- | --- |
| Expression Abundance Threshold | Events Detected | Expression Abundance Threshold | Events Detected |
| Junction coverage > 6 FPKM | 9,277 | Signal > 50% | 10,114 |
| Junction coverage > 2 FPKM | 34,961 | Signal > 25% | 31,506 |
| Junction coverage > 2/3 FPKM | 92,986 | No threshold | 92,405 |
| FDR for significant (p-value < 1e-3) |  |  |  |
| Junction coverage > 6 FPKM | 0.027 % | Signal > 50% | 0.092 % |
| Junction coverage > 2 FPKM | 0.047 % | Signal > 25% | 0.137 % |
| Junction coverage > 2/3 FPKM | 0.070 % | No threshold | 0.345 % |

Table 1 shows that fixing p-value to 0.001 yields False Discovery Rates (FDRs) less than 1% for both technologies. The expected proportion of AS events appears high (1-π_0_ approx. 46 %)[41], i.e. more than 46% of the events have its splicing patterns altered, which reflects the anticipated strong effect of compound exposure on the splicing machinery. It is also apparent that, for a similar number of detected AS events, the FDR corresponding to RNA-seq analysis is smaller.

Events detected by both technologies (referred to as “matched events” hereon) were defined by a stringent criterion in which nucleotide sequences of paths identified via one technology must be a subset of sequences identified via the other, yielding 6,222 matched events. When reporting correspondence and divergence between AS events, below, the following naming convention is used: R^+^ represents number of events deemed significantly altered in RNA-seq analysis; R^−^ represents number of events deemed not significantly altered in RNA-seq analysis. M^+^ and M^−^ are the counterpart terms used to describe microarray results. Events not detected by each technology are labelled R∅ and M∅ respectively.

A subset of matched events is significant in both technologies (R^+^M^+^) and shows coherent change in the corresponding Ψ. There are also significant events detected by only one of the technologies (R^+^M∅ and R∅M^+^). The summary of findings is presented in Table 2.

**Table 2.** Number of AS events detected per technology, alongside statistical significance of events against distinct thresholds. Where thresholds not shown, default filters were employed (junction coverage >2 FPKM for RNA-seq; upper quartile probe signal for microarrays).

| RNA-seq |  | Exon-junction arrays |  |
| --- | --- | --- | --- |
| Condition | Events Detected | Condition | Events Detected |
| Matched events | 6,222 | Matched Events | 6,222 |
| Significant in both ($R^+M^+$) | 1,324 | Significant in both ($R^+M^+$) | 1,324 |
| FDR for matched events RNA-seq | 0.0496% | FDR for matched events arrays | 0.0623% |
| Number of events in $R^+M^\emptyset$ | | Number of events in $R^\emptyset M^+$ | |
| Expression Abundance Threshold | Events Detected | Expression Abundance Threshold | Events Detected |
| Junction coverage > 6FPKM<br>Junction coverage > 2 FPKM<br>Junction coverage > 2/3 FPKM | 2,973<br>10,617<br>25,063 | Signal > 50%<br>Signal > 25%<br>No threshold | 1,016<br>3,297<br>7,581 |
| FDR for $R^+M^\emptyset$ | | FDR for $R^\emptyset M^+$ | |
| Expression Abundance Threshold | FDR | Expression Abundance Threshold | FDR |
| Junction coverage > 6FPKM<br>Junction coverage > 2 FPKM<br>Junction coverage > 2/3 FPKM | 0.0244%<br>0.0456%<br>0.0690% | Signal > 50%<br>Signal > 25%<br>No threshold | 0.146%<br>0.199%<br>0.485% |

Table 2 shows that the FDR of the events detected only by RNA-seq is similar to that for events detected by both platforms (0.0456% vs. 0.0496%). In other words, the reliability of events discovered only by RNA-seq is similar to that of events identified by both technologies. In the case of the arrays, the FDR of matched events is three times smaller than for those discovered solely by the arrays (0.199% vs 0.0623%), i.e. R∅M^+^ events are less reliable than R^+^M^+^ events for the same p-value threshold. In addition, Table 2 shows that the number of significant events that are RNA-seq specific (R^+^M∅) is larger than the number of significant events detected only by arrays (R∅M^+^) (10,617 vs 3,297 events).

Figure 2 depicts a Sankey diagram of the relationship between matched events. It is apparent that many events that are significant for RNA-seq are not detected by arrays, but also that events significantly detected via arrays are not detected by RNAseq. Most matched events are consistent across technologies: significant events for one technology are also significant for the other.

**Figure 2.**
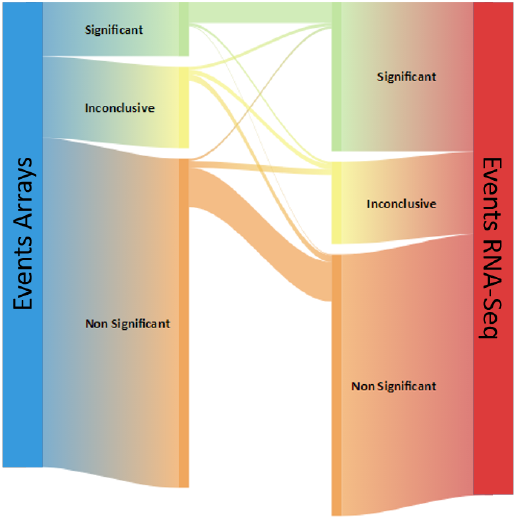
Correspondence between the events detected by arrays and RNA-seq. An event is considered to be significant if the p.value is smaller than 0.001 and non-significant is it is larger than 0.2. Events with p-values between both are considered to be inconclusive cases.

### PCR validation rates are over 80% in both technologies

PCR validation was performed on a subset of predicted AS events drawn from each of the subsets discussed in previous sections, i.e. events detected by one or both technologies. PCRs were performed on:

1. Five top-ranked events detected by both technologies (*topRNA* and *topArrays*) regardless of the matching with the other technology
2. Five top-ranked events detected by one technology (R^+^M∅ and R∅M^+^)
3. Five top-ranked events significant in one technology (R^+^M^−^ and R^−^M^+^)
4. Five top ranked events detected by both technologies (R^+^M^+^)

PCR for events in non-coherent classes (R^+^M^−^, R^−^M^+^) required up to 40 PCR cycles and were harder to validate in general.

A summary of the outcome is shown in

Table 3. The characteristics of validated events (genome location, event type, number of PCR cycles, etc.) are included in Supplementary file 3. The corresponding GTF files to browse these events in IGV[42] are included in Supplementary file 4.

**Table 3.** PCR validation for RNA-seq and microarray technologies across events detected by one or both technologies. Values reported are validations / events selected.

| AS Event Category | RNA-seq | Arrays |
| --- | --- | --- |
| Top-ranked events (*topRNA*, *topArrays*) | 5/5 | 5/5 |
| Significant in RNA-seq and not detected by arrays ($R^+M\emptyset$) | 5/5 | - |
| Significant in arrays and not detected by RNA-seq ($R\emptyset M^+$) | - | 5/5 |
| Detected by both. Significant in RNA-seq, not significant in arrays ($R^+M^-$) | 5/5 | |
| Detected by both. Significant in arrays, not significant in RNA-seq ($R^-M^+$) | 3/5 | |
| Detected by both. Significant and coherent events ($R^+M^+$) | 5/5 | |

Figure 3 shows the Ψ estimates and the PCR bands for two of the top-ranked events in R^+^M^+^ (gene names *DONSON* and *MELK*), with clear concordance of the splice index, Ψ, across the three technologies despite use of end-point (i.e. non quantitative) PCR. Similar figures for events in the other AS categories are included in the supplementary material (Figure S 2 to Figure S 6).

**Figure 3.**
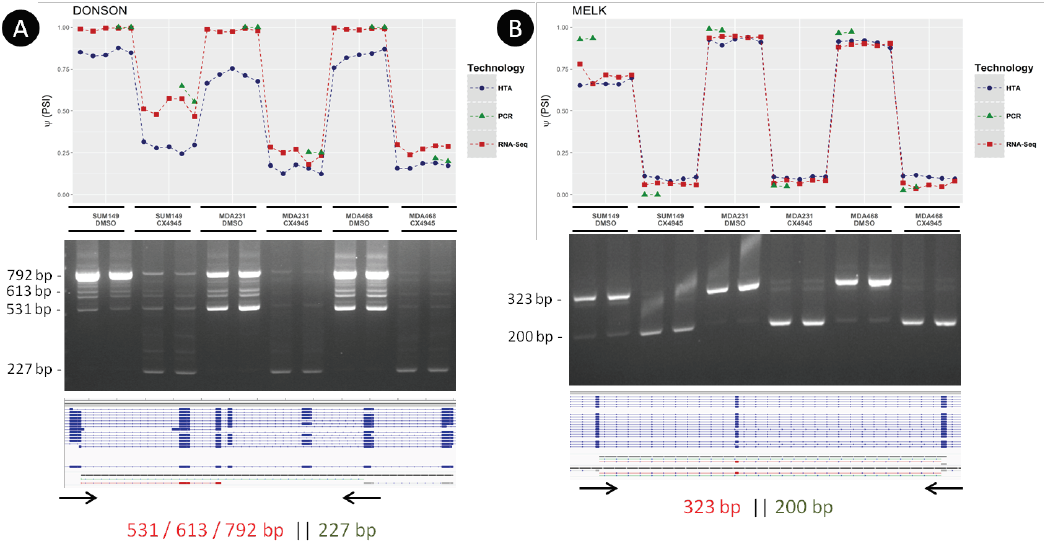
Estimated PSI (for RNA-seq, microarrays and PCR image analysis), PCR bands, the reference HTA transcriptome and the alternative paths of the *DONSON* (panel A) and *MELK* (panel B) gene in *R* ^*+*^ *M* ^*+*^. This concordance occurs for most of the genes validated by PCR. The last numbers shown are expected bands for the selected primers. If the number is shown in red, the band corresponds to Path 1 of the even (long path)t. If shown in green, corresponds to Path 2 (short path).

### Statistics and Ψ for matched events are similar

Figure 4 shows the increment of the Ψ value estimated by EventPointer for events detected by both technologies. Correlation for AS events is over 0.90, and z-values of the statistical test are also similar (Figure S 1). PCR figures also show high coherence between the estimated Ψ using both technologies, especially for RNA-seq, and the PCR results (Figure 3 and Figure S 2 to Figure S 6)

**Figure 4.**
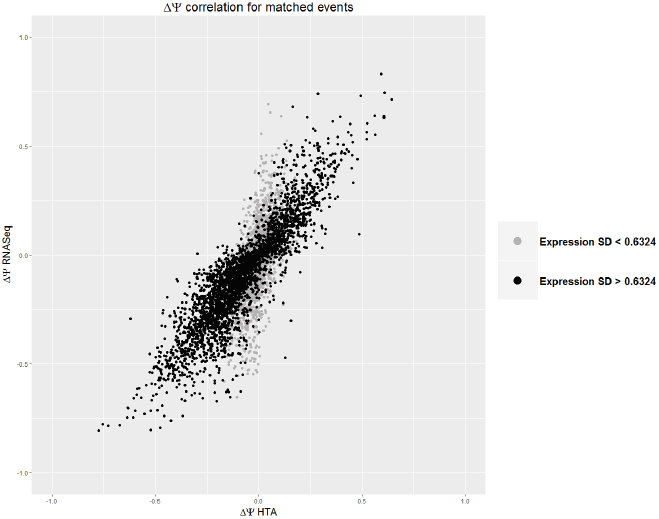
Increment of PSI for both microarrays and RNA-seq. The black (gray) dots represent events with high (low) standard deviation in the differential usage of the isoforms in both paths. Correlation between events with high and low variability are 0.90 and 0.61 respectively.

### Both technologies detect a similar distribution of AS types

Figure 5a shows the number and type of AS events detected by the EventPointer algorithm on data from both profiling technologies. The number of detected cassette exons using arrays is smaller than that using sequencing (p.value < 1e-16, test for equality of proportions). In fact, after matching the events detected by both technologies, a large proportion of the cassette exons in RNA-seq appear as complex in microarrays (see Figure 5b). The reason for this disparity is the complexity of the reference transcriptome used in the HTA array. For this analysis, we used the transcriptome provided by Affymetrix, which includes a range of annotation sources, e.g. RefSeq, Vega, Ensembl, MGC (v10), UCSC known genes and other sources for non-coding isoforms. The underlying transcriptome for HTA includes such a variety of isoforms that many detected AS events are labelled as complex.

**Figure 5.**
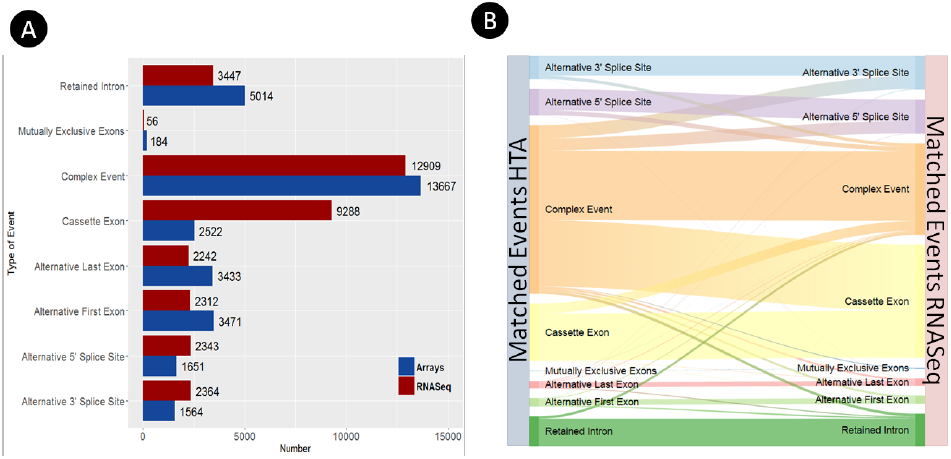
A) Events detected using RNA-seq and array technologies. B) Type of event after matching the events detected by both technologies

In addition, the proportion of retained introns is smaller for RNA-seq (p.value < 1e-16, test for equality of proportions), perhaps owing to the coverage required to include a region as expressed by SGSeq (defaults to 0.5 FPKM) which may exclude weakly expressed introns.

### Power of arrays to detect events is approximately equivalent to shallow RNA-seq

The comparisons above suggest that that RNA-seq – at the depth of sequencing deployed here - detects a larger number of AS events at lower FDR.

Successive approximations using 30% and 10% of the initial RNA-seq reads (approximately 30 million and 10 million reads respectively) yield the estimate that FDR obtained using junction arrays in the present comparison is equivalent to that which would be obtained by an RNA-seq experiment with sequencing depth of approximately 20 million reads.

We hypothesize that the performance of the arrays could be greatly improved by removing bad-performing probes. In fact, we have identified some probes that cross-hybridize in several loci of the transcriptome. Their corresponding events do not show internal coherence with the model (i.e. they show a large relative error if the weighted sum of the signals in Paths 1 and 2 and the reference Path are compared). These bad probes are somehow expected: the design of junction probes has strong limitations since there is no room to select a probe with certain standards of quality (GC content, no cross-hybridization against the genome of the transcriptome, etc.). Owing to these probes, a number of events are not being measured accurately.

The events that are not matched with RNA-seq are enriched in these pathological cases as shown in Table 2. On the contrary, the expected false discovery rate for matched events finds HTA arrays to be equivalent to RNA-seq with a depth of approximately 60 million reads. A proper filtering of the probes identifying events prone to errors could ideally make the arrays equivalent to RNA-seq with this depth.

## Discussion

The main aim of this work was to quantitatively compare the performance of RNA-seq and junction array technologies to detect splicing events. To do this in a balanced manner, we adapted our algorithm EventPointer, originally developed for HTA arrays, to work also on RNA-seq data.

This study highlights:

- The creation of a real-world cell exposure dataset specifically relevant for the study of alternative splicing.
- Adaptation of an existing AS event detection algorithm to a cross-platform method to enable comparative application, and addition of percent splice index method.
- Strong correlation of splicing event detection in regions covered by both technologies, validated by PCR on a subset of top ranked events identified by both and each platform respectively.
- Benefits of RNA-seq in terms of coverage and flexibility, as expected, and higher validation rates in case of disagreement between technologies.
- Good performance of HTA arrays, estimated by approximation to be equivalent to relatively shallow RNA-seq in transcript regions covered.

Top-ranked events detected by each platform technology and estimates of relative event occurrence (ΔΨ) were validated by PCR. The relative occurrence estimates were also strongly correlated, close to 0.90 for events detected by both technologies. In addition to enabling comparison of the two profiling platforms, these results suggest also that the occurrence estimates themselves are a relevant addition to the original EventPointer algorithm.

As might be expected in the absence of physical probe-sets, over 10,000 statistically significant events were identified by RNA-seq alone, the top ranked of which were validated via PCR. Approximately 3,300 events were detected using microarrays but not detected using RNA-seq. In this case, some (3/5) of the top ranked events were validated and correspond to well-expressed genes. Those which did not may reflect the specific technical biases of each technology (cross hybridization of the probes, multi mapping reads, GC dependence, etc.)

RNA-seq has inherent advantages over microarrays, including the ability to detect unlimited novel events. Furthermore, sensitivity can be improved by increasing sequencing depth. Another advantage of RNA-seq is its better approximation of gene/transcript concentrations (e.g. allowing to state a threshold based on the expression of an event). On the other hand, arrays were able to detect some weakly expressed events missed by RNA-seq and, in general across the comparisons, performed similarly to RNA-seq when treating well-expressed and well-defined transcriptional regions. As expected, a similar algorithmic approach applied to both platforms consumed less time and memory resources when treating microarray data than when treating RNA-seq data (See Table 4).

**Table 4.**
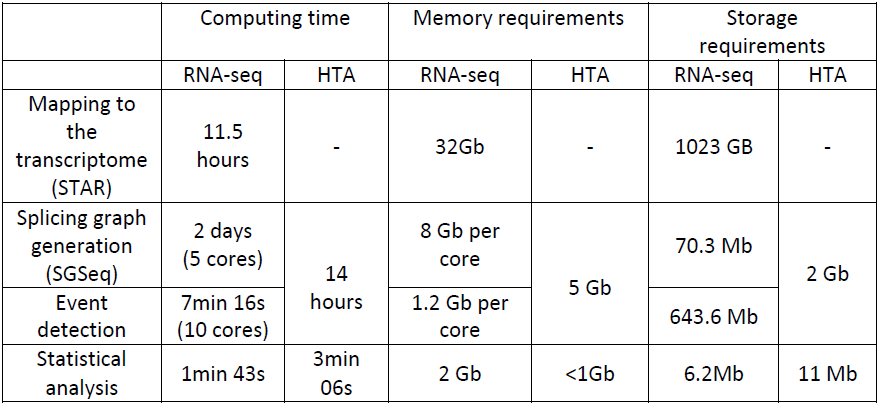
Resources required for both technologies. Analysis was performed on 16 cores (Intel Xeon E5-2670@ 2.60 GHz) with 64 GB of RAM Linux server running 64 bit CentOS distribution

|  | Computing time |  | Memory requirements |  | Storage requirements |  |
| --- | --- | --- | --- | --- | --- | --- |
|  | RNA-seq | HTA | RNA-seq | HTA | RNA-seq | HTA |
| Mapping to the transcriptome (STAR) | 11.5 hours | - | 32Gb | - | 1023 GB | - |
| Splicing graph generation (SGSeq) | 2 days (5 cores) | 14 hours | 8 Gb per core | 5 Gb | 70.3 Mb | 2 Gb |
| Event detection | 7min 16s (10 cores) |  | 1.2 Gb per core |  | 643.6 Mb |  |
| Statistical analysis | 1min 43s | 3min 06s | 2 Gb | <1Gb | 6.2Mb | 11 Mb |

In conclusion, comparison of RNA-seq and junction microarrays using a cross-platform algorithm suggests that both technologies provide accurate identification of splice events. Moreover, predictions by both technologies tend to correlate strongly and yield similar results when compared by Ψ estimates and PCR. RNA-seq holds a clear advantage in terms of flexibility, and stronger PCR validation of events detected in one platform but not the other. As compared, HTA microarrays are shown nevertheless to provide a reasonable alternative to relatively shallow RNA-seq in the transcriptional regions that they reference.

## Methods

### Sample preparation

Triple negative breast cancer cell lines MDA-MB-231 and MDA-MB-468 were obtained from ATCC (Manassas, VA) and SUM149 was purchased from Asterand plc (Detroit, MI). All cell lines were grown according to the suppliers’ recommendation. CK2 inhibitor CX-4945 (Selleckchem, Houston, TX) was dissolved in DMSO and stored frozen at −80°C until used.

To induce splicing events, cells were grown to ˜70% confluence and treated with 1µM CX-4945 or DMSO during 12 hours in a total of 5 replicates per condition. Total RNAs were isolated using the RNeasy Mini Kit (Qiagen, Germantown, MD) according to the manufacturer’s protocol. Integrity of RNA was quantified using the Agilent 2100 Bioanalyzer (Agilent Biosystems, Foster City, CA). Samples were labeled and hybridized in Human Transcriptome arrays (HTA) by the Genomics Core Facility of the Center for Applied Medical Research (CIMA) following manufacturer’s instructions.

RNAseq was performed in the Center for Cooperative Research in Biosciences (CICBiogune) using the Illumina HiSeq2000 sequencing technology, HiSeq Flow Cell v3 and TruSeq SBS Kit v3. 2ug of RNA of each sample was sent for this purpose. The run type was strand specific, multiplexed with paired-end reads of 100 nucleotides each. The amount of RNA for hybridization and validation purposes was 5 ug.

STAR 2.4.0h1 was used to align the reads against the human genome. The reference genome was Ensembl GRCh37.75. The output were sorted BAM files. All the other parameters were set to the default values. The average sequencing depth was 49 million reads (9.8 billion nucleotides sequenced per sample).

The microarray data preprocessing was performed using the aroma.affymetrix framework using the standard RMA algorithm[36].

### Event Pointer for RNAseq

EventPointer is an R package to identify, classify and analyze alternative splicing events using microarrays and RNA-Seq data. The software is available for download at Bioconductor. A thorough description of EventPointer for microarrays can be found in [8]. This method has been extended to RNA-seq.

The concepts for detection, classification and statistical analysis are shared in EventPointer for the analysis of both technologies. The main difference of EventPointer for RNA-seq compared with that of microarrays are the ones associated with the type of input data (CEL or BAM files).

EventPointer requires a splicing graph -a directed graph used to represent the structure of the different isoforms of a given gene[43]- as input to detect splicing events. EventPointer for RNA-seq uses SGSeq[30] to build the corresponding splicing graphs from BAM files. The complexity of the splicing graph can be controlled in SGSeq by setting different thresholds in the expression values of splicing junctions of the splicing graph (by default set to 2 FPKM). For RNA-seq, the splicing graphs are constructed for every single experiment. On the contrary, in the case of microarrays the same splicing graph (and the corresponding CDF) is used for all the experiments run on the same type of microarray (HTA or, more recently Clariom-D).

The input data for the statistical analysis is different in both technologies: signal values of the probes in microarrays and counts in RNA-seq. In order to deal with reads, Voom[16] is applied to preprocess the RNA-seq count data. The statistics to deal with the processed RNA-seq data is identical to the one used for microarray data and hence, the same statistical tests -based on limma[39]- are applied to both technologies.

As output, EventPointer provides a table with the following information associated to each detected alternatively spliced event: gene identifier, genomic position, type of event, statistical parameters and ΔΨ values. Additionally, EventPointer generates a “Gene Transfer Format” (GTF) file that can be used with the Integrative Genomics Viewer (IGV)[42] to view the structures of each detected alternative splicing event. This visualization facilitates the interpretation of the detected events and the design of primers for the validation of the events using standard PCR.

### Estimation of PSI

We have included a novel algorithm to estimate Ψ that can be applied to both RNA-seq and microarrays. Assuming that the signal of a probe-set in microarrays and the number of reads within a region of the transcriptome in RNA-Seq depend on the product of an affinity value of the probe-set (or the equivalent length in RNA-seq) and the concentration of the interrogated isoforms in the paths, the following equation holds

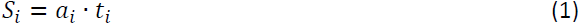

where *S*_*i*_ is the measured expression value of path *i*, *a*_*i*_ is the affinity of the probes or equivalent length of the path *i* and *t*_*i*_ is the concentration of the isoforms mapped to path *i*. The affinity values (or equivalent lengths) and concentration values are unknown and must be estimated from the data.

Particularizing the above equation to each of the paths and taking into account that the concentration of the isoforms in the reference path must be the sum of those of paths 1 and 2, the following equations are obtained:

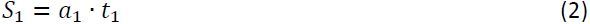

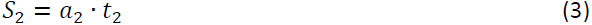

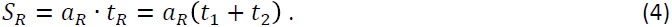

In turn, the signal value of the reference path can be expressed as the sum of the signal values of paths 1 and 2 as follows,

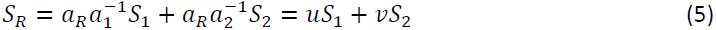

where *u* and *v* represent the fraction of the affinities of the mapped probe-set (or equivalent lengths) in the reference path and paths 1 or 2 respectively. The values of *u* and *v* can be estimated from signal data.

Dividing equation (2) with equation (4) we get,

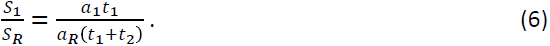

Combining equations (5) and (6), the desired equation of the Percent Spliced Index (Ψ) used in EventPointer is obtained:

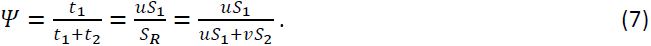

Note that Ψ can be directly obtained from signal values once *u* and *v* are known. This equation does not require the estimation of the affinities (difficult to predict accurately) to compute Ψ. On the contrary, it simply requires to estimate *u* and *v* from signal values using equation (5). In the case of RNA-seq, the equivalent lengths are known a priori and hence *u* and *v*. However, using this approach has an advantage: the estimates of these lengths can accommodate the potential lack of uniformity of the reads.

Note that *u* and *v* must be positive, similar between them and close to one. The first affirmation is trivial since affinity values (or equivalent lengths) are always positive. In microarrays, probe-sets are composed by several probes and their overall affinity are expected to be similar to each other, since these affinities are a median of the average of the affinities of the probes that build up them. Therefore *a*_1_ ≈ *a*_2_ ≈ *a*_3_, and *u* ≈ *v* ≈ 1. A similar reasoning can be applied to RNA-seq,if using coverage instead of read coutns, since the coverage of the reference path is expected to be close to the sum of the coverages of paths 1 and 2.

These two fractions can be estimated from equation (5) by using non-negative least squares as follows:

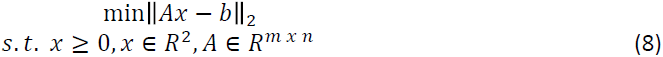

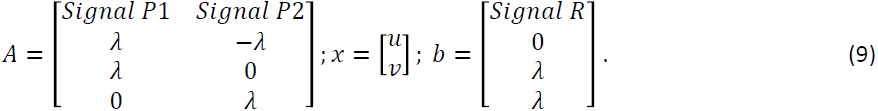

The penalty factor λ is added to force the equation to fulfill the previous considerations: *u* and *v* must be similar and close to 1. In our results, we found that the estimates were not too sensitive to the specific value of λ if there is differential alternative splicing. If the relative usage of both paths is similar and therefore, Ψ is constant, the results are more sensitive to the value of Ψ. This fact is shown in Figure 4: the correlation is much better for events that show variability in the relative expression of both paths.

### Statistical analysis

The comparison and analysis of the profiling data was done using a linear model. The design matrix was built considering both the cell line and treatment with CX4945 as factors. The interaction between cell line type and treatment was not considered.

The selected contrasts test for the difference between control samples (DMSO) and drug exposed ones (CX4945) controlling for the cell-type. The complete experimental design in the form of design and contrast matrices is included in Table Supplementary file 5.

EventPointer includes several statistical methods to state the significance of an event. In this experiment, the events are considered to be statistically significant if there is a change in the expression of the isoforms associated to each of the alternative paths, this change occurs in, opposite direction, i.e. opposite signs for the fold changes and the summarized p.value is significant (p.value< 0.001).

### Filters used to include the events

For arrays, the signal of the probe-sets interrogating each of the alternative paths involved in a splicing event, must be expressed more than a certain threshold in at least one sample. This threshold is the 25% quantile of the expression of the signal in the reference paths for all the events included in the array. For RNAseq, the edges of the splicing graph (junction reads) are included only if their expression is at least 2 FPKM in at least one sample (SGSeq defaults).

### Matching of the events using different technologies

Let’s assume that *A*_*R*_ and *A*_*M*_ are, possibly non-contiguous, regions of the genome that correspond to path A using either technology (*A*_*R*_ for RNA-seq and *A*_*M*_ for HTA). *B*_*R*_ and *B*_*M*_ have a similar description for path B and *R*_*R*_ and *R*_*M*_ for the reference path in each technology. Two events are considered to match if any of the following two expressions is true:

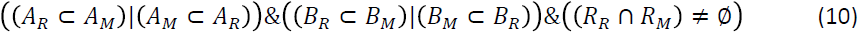

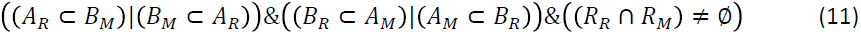

In these expressions, (*x* ⊂ *y*) is true if the genomic region *x* is a subset of the genomic region *y* (the nucleotide sequence of *x* is a substring of the nucleotide sequence in *y*). Besides, the operators “|” and “&” and the logical OR and AND operations. If (*x* ⊂ *y*)|(*y* ⊂ *x*), then one of the regions is contained in the other are considered to be “compatible". On the other hand,(*x* ∩ *y*)≠ ∅ means that regions x and y overlap in the genome. Therefore, the first expression is true if both paths *A*_*R*_ and *A*_*H*_ are compatible, *B*_*R*_ and *B*_*M*_ are compatible and *R_R_* and *R_M_* overlap. The second expression is true if path *A*_*R*_ and path *B*_*M*_ are compatible and also path *A*_*M*_ and *B*_*R*_ are compatible and, again, and *R_R_* and *R_M_* overlap.

Within an event, the longer path in the transcriptome is assigned the name “A” and the other the B. The second equation (11) takes into account that, in some few cases, the name of the paths can be switched in both technologies.

### PCR validation

For each splicing event, an end-point PCR was run using primers designed in the exons that flank the event of interest. RNA was retro-transcribed and the PCR was performed an analyzed as previously described[44]. Primers used are shown in Table Supplementary file 3.

## Data Access

EventPointer for both RNA-seq and microarrays is available at Bioconductor.

All the RNA-seq and microarray data are available in a superseries at Gene Expression Omnibus, access number GSExxxx.

## Acknowledgements

The work performed and described was funded by Celgene Research SL, part of Celgene Corporation. Author FC was partially supported by a Basque Government predoctoral Grant [PRE_2016_1_0194]. LMM and RP were partially funded by Spanish Ministry of Economy and Innovation and Fondo de Investigación Sanitaria-Fondo Europeo de Desarrollo Regional (PI14/00806, PI16/01821), CIBERONC and AECC Scientific Foundation (GCB14-2170).

## Author Contributions

**Conception and design:** J.P. Romero, M. Ortiz-Estévez, M. Trotter, A. Rubio

**Development of methodology:** J.P. Romero, A. Rubio, A. Muniategui

**Acquisition of data (provided cell-lines treatment, provided sequencing and hybridazation, performed PCR validations, provided facilities, etc.):** S. Carrancio, F.J. de Miguel, R. Pio, L.M. Montuenga, M. Trotter

**Analysis and interpretation of data (e.g., statistical analysis, biostatistics, computational analysis):** J.P. Romero, A. Rubio, R. Loos, M. Ortiz-Estevez

**Writing, review, and/or revision of the manuscript:** J. P. Romero, M. Ortiz-Estévez, A. Muniategui, L.M. Montuenga, R. Loos, R. Pío, A. Rubio, M. Trotter

**Administrative, technical, or material support (i.e., reporting or organizing data, constructing vignettes):** F. Carazo, J.P. Romero, A. Muniategui, A. Rubio

**Study supervision** R. Loos, A. Rubio, M. Trotter

## Footnotes

1 Average sequencing depth for RNA-seq was approx. 98 million (paired-end, stranded protocol), yielding on average approx. 49 million fragments per sample.

